# Spatial Distribution of Cortical Output Zones Affecting Combinations of Forelimb Muscles in the Monkey

**DOI:** 10.64898/2026.06.24.731406

**Authors:** Paul D. Cheney, Suzanne Sawyer Vincent, Richard F. Martin, Eberhard E. Fetz

## Abstract

**SUMMARY AND CONCLUSIONS:**

1. We investigated the dimensions of output zones affecting specific combinations of forelimb muscles in the precentral “motor” cortex of macaque monkeys. Single-pulse intracortical microstimulation (S-ICMS) was used to evoke subthreshold effects in multiple wrist and finger muscles. S-ICMS consisted of 15 Hz stimulus trains at low intensity (5 μA) to avoid spread of effects due to temporal summation and to activate circumscribed foci of cortical cells in the bank of the precentral gyrus.
2. To detect subthreshold effects on active muscles, stimuli were delivered during wrist movements against elastic loads, and stimulus-triggered averages of rectified electromyographic (EMG) activity of 12 identified forelimb muscles were compiled. These averages detected statistical increases and decreases in averaged rectified EMG activity termed poststimulus facilitation and poststimulus suppression, respectively (7, 8). The “muscle profile” of a cortical output site was defined as the distribution and relative magnitude of these effects evoked in the recorded muscles. To document the spatial extent of a cortical output zone producing a particular “muscle profile,” we delivered S-ICMS at successive sites along electrode tracks in the precentral bank, tangential to and near layer V cells.
3. In some cases the electrodes encountered cortical cells whose post-spike effects on muscles were also documented by spike-triggered averages of EMG activity. Near cells that had post-spike effects, S-ICMS evoked a similar profile of effects on these muscles, over distances of several hundred microns from the location of the cells with post-spike output effects.
4. The locations of tracks were marked by electrolytic lesions at specific depths and were identified in subsequent histological reconstructions. The physiological effects evoked from different sites were correlated with the locations of Nissl-stained cortical cells and corticospinal cells labeled by horseradish peroxidase (HRP) transported from the cervical spinal cord.
5. The “muscle profiles” of poststimulus effects elicited by 5-10 μA S-ICMS applied at successive sites were the same for multiple neighboring sites extending over tangential distances of about 1 mm. S-ICMS applied at 300 to 500 μm intervals evoked significant poststimulus facilitation or suppression effects over a range of 650-1250 μm. Assuming an effective excitation radius of 65 μm for 5-μA current pulses, the lower limit of the mean tangential dimension of cortical output zones on the precentral bank (n = 17) was estimated to be 800 μm.
6. A specific muscle could be affected, either singly or in combination with synergist muscles, from aggregate regions of output sites extending about 5 x 4 mm mediolateral x dorsoventral. These regions were comparable in size to output areas containing all the corticomotoneuronal (CM) cells that produced postspike facilitation in that muscle.
7. The cortical entry points of tracks with CM cells producing postspike facilitation in specific muscles were reconstructed for three monkeys. In two monkeys, the flexor and extensor CM cells were thoroughly intermingled; in the third, the flexor CM cells were preferentially located medially and showed less overlap with the region containing extensor CM cells.
8. These results indicate that each motor cortex site represents a different combination of muscles. The effects evoked from cortical sites separated by several hundred microns invariably involved different profiles of muscle activity. The muscle fields of remote CM cells were rarely identical, while the fields of neighboring CM cells were often similar. Given the number of unrecorded muscles, we conclude that primate motor cortex is a mosaic of output sites representing forelimb muscles in different combinations.

## INTRODUCTION

The spatial organization of output effects evoked from the motor cortex of nonhuman primates has been investigated repeatedly since Ferrier (11) first mapped the movements produced by faradic stimulation of different cortical sites. The evolution of these studies has recently been reviewed (24). Stimulating the surface of the precentral gyrus, investigators have produced maps of the threshold movements evoked in different joints and contractions of specific muscles (6). These maps have represented the threshold excitatory effects evoked and have rarely included measures of inhibition. For example, Chang et al. (6) documented the tension produced in eight muscles of the ankle joint during systematic surface stimulation of motor cortex; some sites produced “solitary responses” in specific muscles while others produced co-contractions. These observations were interpreted in terms of overlapping mosaics of cells that affected individual muscles, although the authors acknowledged that stimuli could subliminally affect additional muscles.

Excitatory postsynaptic potentials (EPSPs) evoked in cervical motoneurons by anodal surface stimulation showed that “colonies” of corticomotoneuronal (CM) cells projecting to single forelimb motoneurons were distributed over cortical regions of 1 x 3.5 to 6 x 3 mm mediolateral x rostrocaudal (22, 28B). Detailed mapping with minimal surface stimulation indicated that colonies of single hindlimb motoneurons on the convexity of area 4 are large (3-7 mm^2^) and can occupy separate locations. The colonies of motoneurons of different muscles overlapped extensively.

Going beyond surface stimulation, intracortical microstimulation (ICMS) (3, 4, 13, 16, 21, 28, 32) allows the exploration of output effects evoked from the depths of the motor cortex. Again, the most convenient measures of output effects have been overt movements or electromyographic (EMG) activity, elicited in a resting or anesthetized animal by high-frequency (200-500 Hz) stimulus trains. Repetitive ICMS is necessary to generate sufficient temporal summation at cortical and spinal levels to elicit contraction. Using repetitive ICMS to determine the topographic organization of output zones to hand muscles from the motor cortex of capuchin monkey, Asanuma and Rosen found that a particular response, e.g., thumb extension, activated by trains of 5- to 10-μA pulses, was obtained from regions on the bank that measured 1 x 8 to 3 x 3 mm dorsoventral x mediolateral (3). These regions included several discontinuous “efferent zones” for each muscle, and overlapped with regions to different thumb, digit, or wrist muscles. The efferent zones had widths of about 800 μm. The sites from which particular movements could be evoked by repetitive stimulation in conscious macaques were also widely distributed (21). Intracortical mapping of the precentral bank of the baboon demonstrated that cortical “aggregations,” composed of colonies of cortical cells projecting to motoneurons of the same muscle, were as large as 6 x 6 mm dorsoventral x mediolateral (1). Aggregations affecting different muscles overlapped substantially.

As a means of mapping motor cortex output sites, the technique of evoking overt responses by repetitive stimulation has two significant limitations. Overt muscle activity represents only the “tip of the iceberg,” and does not reflect either subthreshold excitatory effects or possible inhibitory effects. These additional effects are part of the muscle synergies represented at a site. Second, by temporal summation repetitive stimuli can recruit widespread effects that may be mediated by indirect synaptic connections (17); these could include axon reflexes via fibers arising from remote cells. Indeed, in comparative studies, repetitive ICMS at many sites stirred up responses in muscles that were not affected or were inhibited by single stimuli (16). These limitations are largely resolved by using S-ICMS in combination with stimulus-triggered averaging (stimulus-TA) of EMG in multiple muscles. Stimulus-TAs of muscle activity can detect subthreshold poststimulus facilitation and suppression of the muscles’ motor units; simultaneous stimulus-TAs of multiple muscles can reveal the distribution of these effects in many flexor and extensor muscles. Moreover, these short-latency effects are produced as a direct consequence of a *single* cortical stimulus and therefore more likely to represent the output of the site.

Even S-ICMS is likely to evoke a descending volley in a potentially diverse population of corticospinal cells. The ultimate level of resolution is the output effect of a single corticospinal cell, which can be documented by the post-spike effects produced in spike-triggered averages (STAs) of EMG (12). STAs have shown that single corticomotoneuronal (CM) cells can produce post-spike facilitation in multiple target muscles (on average about 40% of the recorded synergist muscles). These divergent effects are consistent with antidromic stimulation of pyramidal tract neurons (PTNs) from multiple motoneuron pools (4) and divergent patterns of terminal branches of corticospinal cells labeled with horseradish peroxidase (HRP) (31). Thus, even single CM cells can represent a group of synergist muscles as their output targets. In addition, many CM cells also produce post-spike suppression of antagonists of their targe muscles.

In previous experiments, S-ICMS delivered at the site of CM cells produced a profile of poststimulus facilitation of EMG across different muscles that usually matched the profile of postspike facilitation of the CM cell (7, 8). The poststimulus effect of S-ICMS as low as 5 μA was about six times greater, suggesting that S-ICMS activated a group of cells that had similar effects.

Our purpose in the present experiments was to further investigate the nature and spatial dimensions of cortical output zones to wrist and digit forelimb muscles. We delivered low intensity S-ICMS at successive cortical sites in penetrations of the precentral bank of motor cortex and documented the subthreshold poststimulus effects on multiple active muscles. The stimulus sites were related to the location of CM cells and correlated with the location of Nissl-stained cortical neurons or HRP-labeled corticospinal neurons. A preliminary report of these results has been presented (30).

## METHODS

### Recording

Data was obtained from four rhesus macaques (Macaca mulatta) using techniques previously described (7). A 20-mm diameter recording chamber over the precentral arm area (4 mm anterior to bregma, 18 mm lateral) held a remotely controlled hydraulic microdrive to advance tungsten microelectrodes into the cortex.

EMG activity of 12 forearm muscles was recorded differentially from pairs of transcutaneously inserted multistranded stainless steel wires (AS 632 Bioflex insulated wire, Cooner Sales Co., Chatsworth, CA). The EMG wires were taped along the arm and led to a terminal connector at the upper arm. Forearm muscles were identified by anatomic location and responses to intramuscular stimulation, as described previously (7, 8). Muscles recorded were: extensor digitorum two and three (ED2,3), extensor digitorum four and five (ED4,5), extensor carpi ulnaris (ECU), extensor digitorum communis (EDC), extensor carpi radialis brevis (ECR-B), extensor carpi radialis longus (ECR-L), flexor digitorum sublimis (FDS), flexor digitorum profundus (FDP), flexor carpi ulnaris (FCU), palmaris longus (PL), flexor carpi radialis (FCR), and pronator teres (PT).

### Training

The monkeys sat inside a sound-attenuating booth and performed ramp-and-hold wrist movements alternating between flexion and extension zones. Movements were made against elastic loads opposing both flexion and extension directions with zero load at a midpoint with the wrist in a neutral position. Correct responses were reinforced with a small amount of apple- or banana-sauce after every one to three successful alternating movements. A successful movement involved holding the hand in a 10° target zone for 1 s at an angular position of 22 ° in the flexor or extensor directions; this required a static torque of about 106 dyne-cm. Daily recording sessions usually lasted 4-6 hours. The behavioral paradigm has been described in more detail previously (7).

### Data analysis

A preliminary survey of motor effects elicited from intracortical sites was made using repetitive high-frequency ICMS (20- to 50-ms train of biphasic 0.2-ms pulses at 400 Hz). In selected tracks, stimulus-TAs of muscle activity were computed for microstimuli delivered at low intensity (5-10 μA) and low frequency (15 pulses/s) during the extension or flexion hold phase. At 15 pulses/s the interval between stimuli was 67 ms, too long for temporal summation of evoked effects. The S-ICMS pulses triggered an average of full-wave rectified EMG activity beginning 5 ms before and ending 25 ms after the S-ICMS pulses, digitized at 4 kHz. Most stimulus-TAs included 500 triggered events.

Spike-TAs from cortical cells related to wrist movements also were compiled. These cells usually were located at the sites of strongest post-stimulus effects. Post-spike effects were characterized as strong, moderate or weak based on mean baseline-to-peak amplitude (8). The weak postspike effects were excluded from analysis to avoid possible contamination with noise.

To detect any possible crosstalk between recordings from different muscles we computed EMG-triggered averages (EMG-TAs) of agonist muscle EMG activity triggered from motor unit potentials of an individual muscle. All EMG recordings used in this paper had less than 10% crosstalk (for further discussion, see ref. 7).

Some electrode tracks down the precentral bank of area 4 were oriented to run tangential to layer V cells. Neuronal activity was recorded for distances of over 4000 μm in such tracks. S-ICMS was applied every 300-500 μm to determine the tangential dimension of specific output zones. To document position a lesion usually was placed at a site deep in the track by applying 20 μA of anodal DC current for 20 s. Electrode penetrations were made systematically along the bank mediolaterally every 500- 1000 μm.

The amplitude of poststimulus effects, defined as the peak minus the baseline, was plotted as a function of the cortical depth of stimulus sites. Above and below the site from which the maximal effect was obtained, the amplitude of the effect usually decreased continuously. From such depth-amplitude plots, we determined distances over which poststimulus effects in individual muscles were greater than half the maximum peak amplitude (rounded off to the nearest 50 μm). This distance, called the half-width, was used to compare the dimensions of different output zones.

In addition, the poststimulus effects were statistically compared with maximum baseline noise fluctuations, computed by averaging the three maximum baseline fluctuations in the first 10 ms of stimulus-TAs compiled at four different sites in a single electrode track. These fluctuations occurred 5 ms before and 5 ms after the trigger, i.e., prior to any induced poststimulus effects, which began at latencies greater than 5 ms.

The amplitude of the peak above baseline was compared with the amplitude of these noise fluctuations by the t-test. Distances over which poststimulus effects were statistically highly significant (P < 0.01) were determined from the depth-amplitude graphs, rounded off to the nearest 50 μm, and tabulated. The dorsoventral dimension of cortical output zones was estimated from these statistical determinations. The peak:noise ratio was calculated by dividing the amplitude of the poststimulus effect by the mean noise-to-baseline fluctuations.

The microstimulus pulses, EMG activity of 12 muscles, and wrist torque and position were recorded on a multiplexed 16-channel Honeywell 7600 tape recorder (3-kHz bandwidth).

### Histology and reconstruction of maps

At the end of the recording sessions, the monkeys were anesthetized with pentobarbital and perfused with physiological saline followed by 10% formalin. The cortical region under the chamber was blocked and removed. In three monkeys, parasagittal sections of the cortex were cut at 80 μm and stained with cresyl violet Nissl stain. In one monkey (“OS”), corticospinal cells were labeled by injecting 20 μl of 33% HRP in phosphate buffer into the cervical spinal cord from C5 to T2 on the side contralateral to the recording chamber. After 72 h, the animal was perfused with 1% paraformaldehyde and 2.5% gluteraldehyde in phosphate buffer. The HRP- labeled corticospinal cells were reacted with tretramethyl benzidine (TMB) and counterstained with neutral red Nissl.

Before removing the tissue block, the cortical surface was photographed to obtain an image that could be projected onto a map of the electrode tracks constructed from chamber coordinates. Fig. 9 illustrates a systematic series of electrode tracks made in monkey OS. The locations of stimulus sites and HRP-filled corticospinal cells were visualized in the original histological parasagittal sections (dashed lines) and projected into reconstructed planes perpendicular to the central fissure (solid lines). Parasagittal sections are shown in Figs. 3, 7 and 8 and reconstructed planes in Figs. 10 and 11. To represent the locations of activated corticospinal cells in layer V a flattened map was also reconstructed by projecting the stimulus sites in white matter to a corresponding layer V site, guided by the orientation of labeled axons and then rendering a flat plane of layer V. This showed the distribution of sites from which specific muscles could be affected (e.g. Figs. 11–13 and supplemental figures S1-S2).

**FIG. 1.**
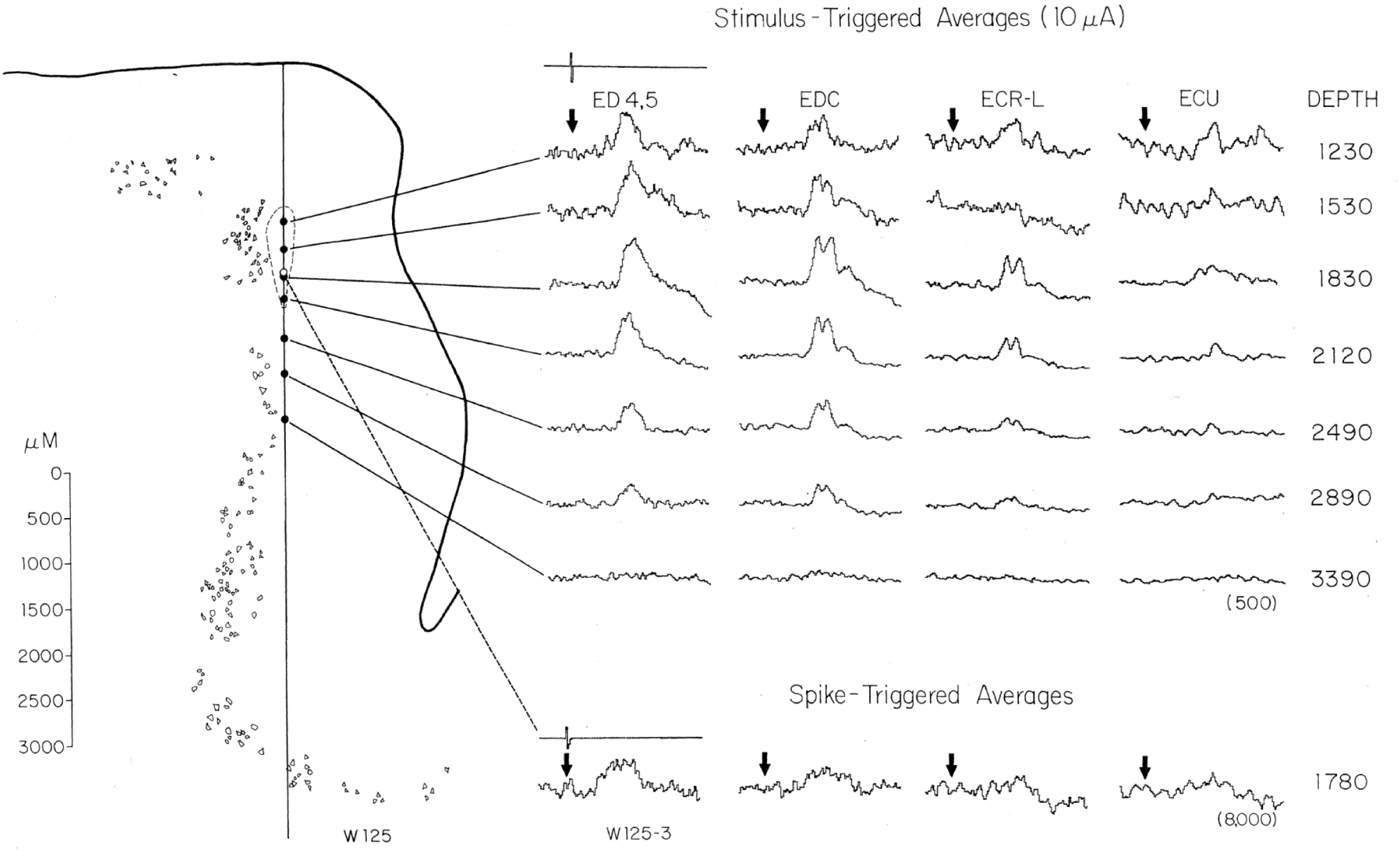
Poststimulus facilitation of extensors in a track tangent to layer 5. Poststimulus facilitation matched the profile of postspike facilitation obtained from cell W125-3 (open circle and bottom traces). Filled circles mark the sites stimulated with 10-μA S-ICMS. Location of Nissl-stained layer V cells on the section are shown schematically. The lesion placed at depth 1619 is outlined by dashes. Where tested (from depth 600 to 1449), reciprocal poststimulus suppression was evoked in FCU and PL (not shown). In this and succeeding figures the numbers in parentheses indicate numbers of samples in the averages. The duration of averages was 30 ms: 5 ms before and 25 ms after trigger (arrows).

**FIG. 2.**
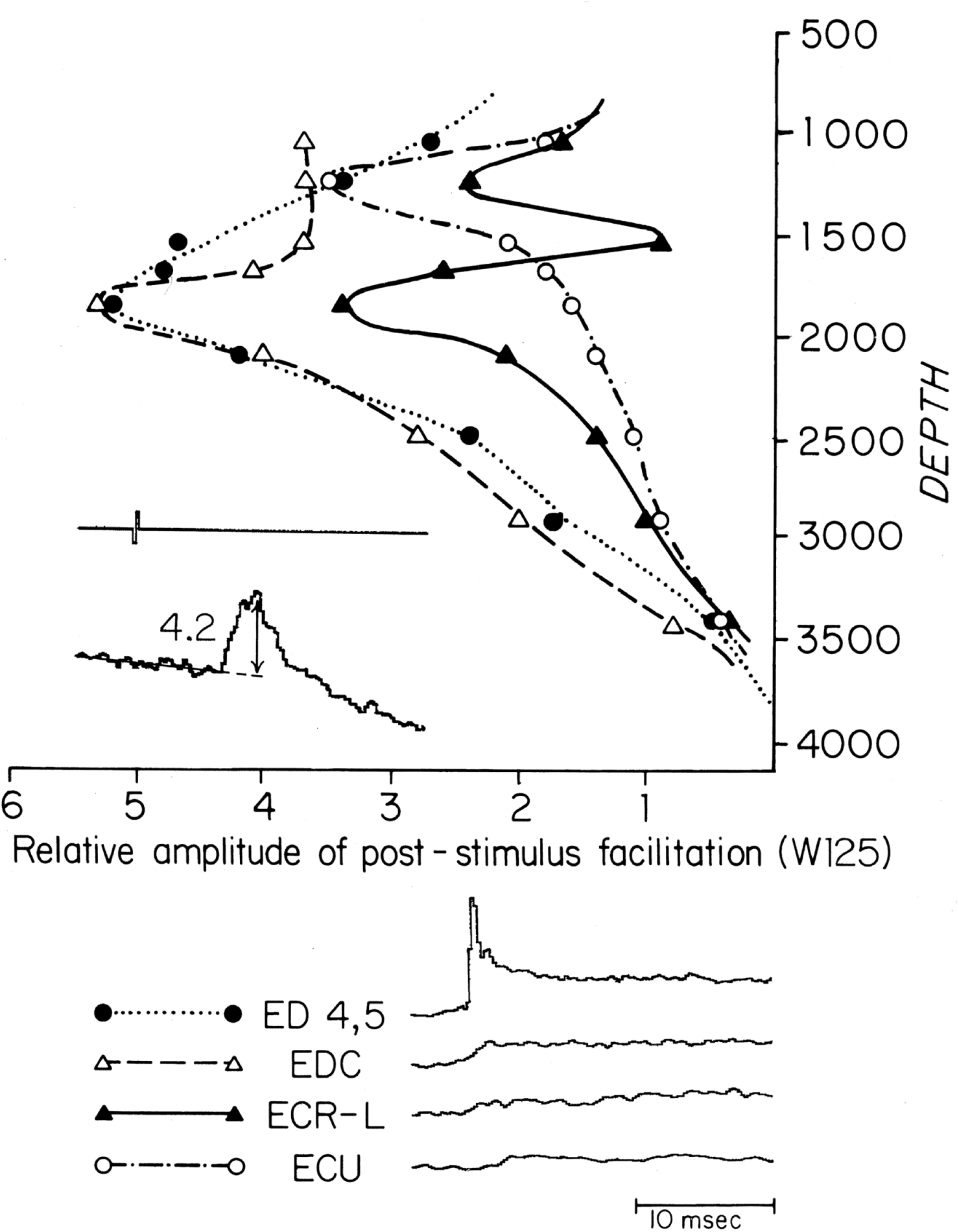
Top: amplitude of poststimulus facilitation plotted as a function of depth of stimulus site in track W125 (cf. Fig. 1). Amplitude of poststimulus facilitation (evoked by 10 μA) was measured baseline-to-peak, as indicated by the inset; the relative amplitude illustrated was 4.2 in arbitrary units. Effect on ED4,5, EDC and ECR-L peaked at depth 1800; peak effect on ECU was from a more superficial site, depth 1200. Bottom: EMG-TAs of extensor muscles recorded in this experiment. Motor unit potentials of ED4,5 were used to trigger averages of rectified EMG activity in EDC, ECR-L and ECU. ED4,5 showed a peak at time 0 reflecting summation of triggering motor unit potentials. No significant crosstalk between ED4,5 and other extensor muscles was observed (7).

**FIG. 3.**
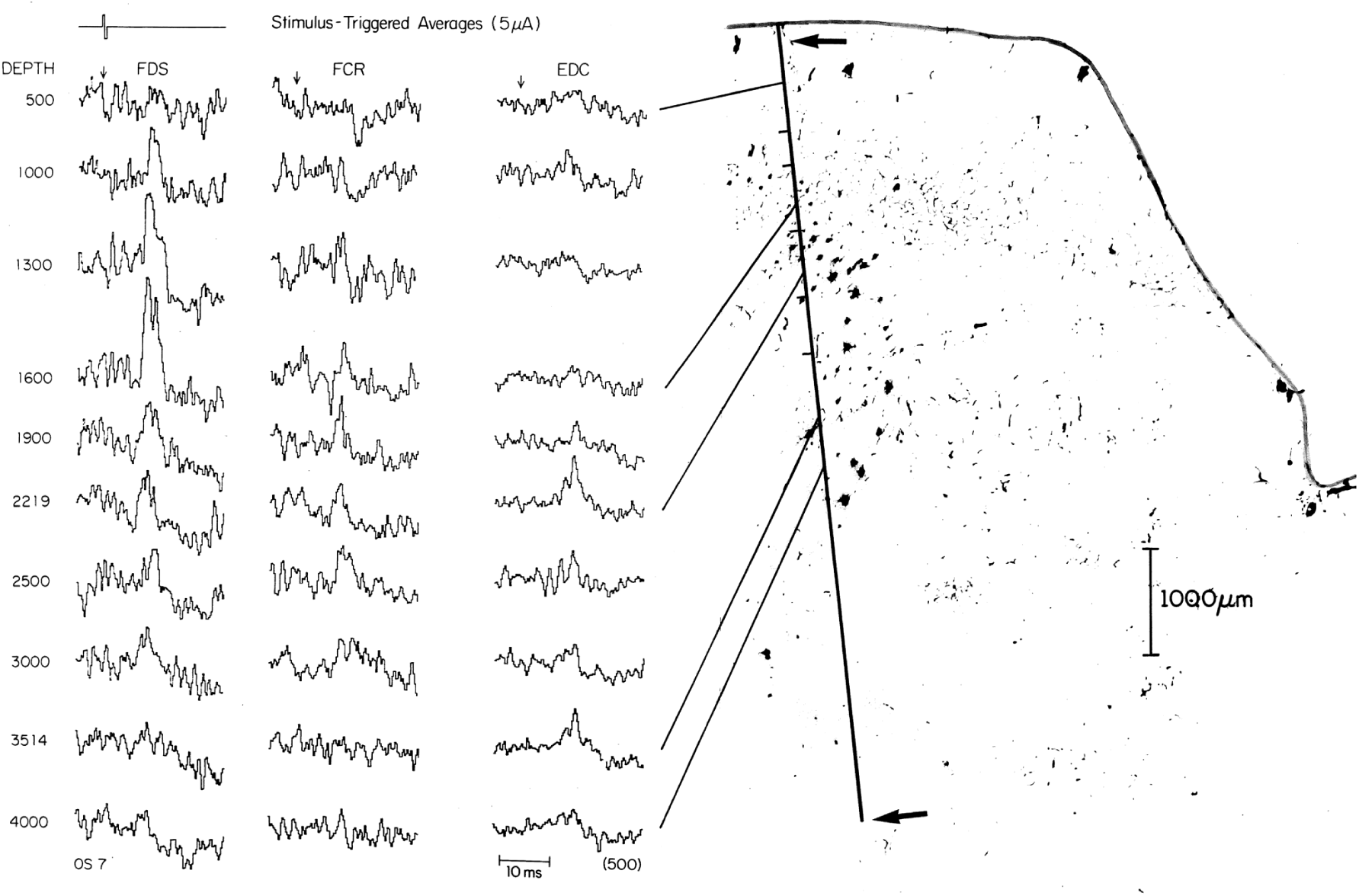
Histologic section containing track OS7 and corresponding stimulus- TAs. Arrows indicate entry point and lesion at depth 7200. Averages aligned by depth are indicated by either a solid line connecting averages to the electrode track or by a short line to the left of the track. Dark cells are corticospinal neurons labeled by retrograde transport of HRP from cervical cord. FDS showed facilitation at superficial sites in layer V (maximal at depth 1600); EDC exhibited maximal poststimulus facilitation at deeper sites (depths 2219 and 3514). FCR showed more complex effects including a mixture of poststimulus facilitation and suppression.

**FIG. 4.**
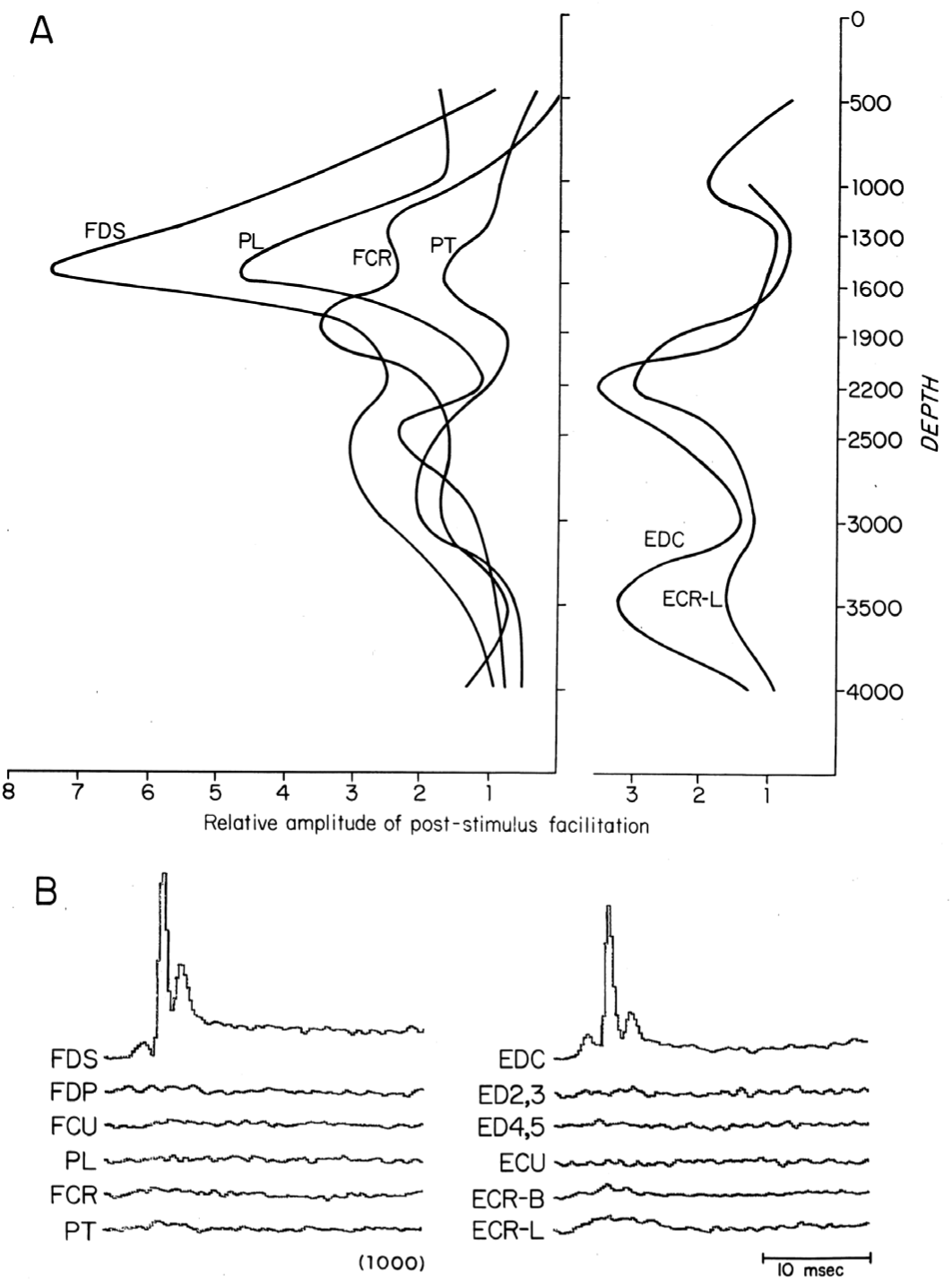
A: amplitude of poststimulus facilitation versus depth in flexor and extensor muscles evoked from microstimulation in track OS7 (cf. Fig. 3). B: EMG-TAs of flexor muscles triggered from FDS and extensor muscles triggered from EDC confirmed that effects in multiple muscles, e.g., FDS and PL, were not due to crosstalk between EMG recordings.

**FIG. 5.**
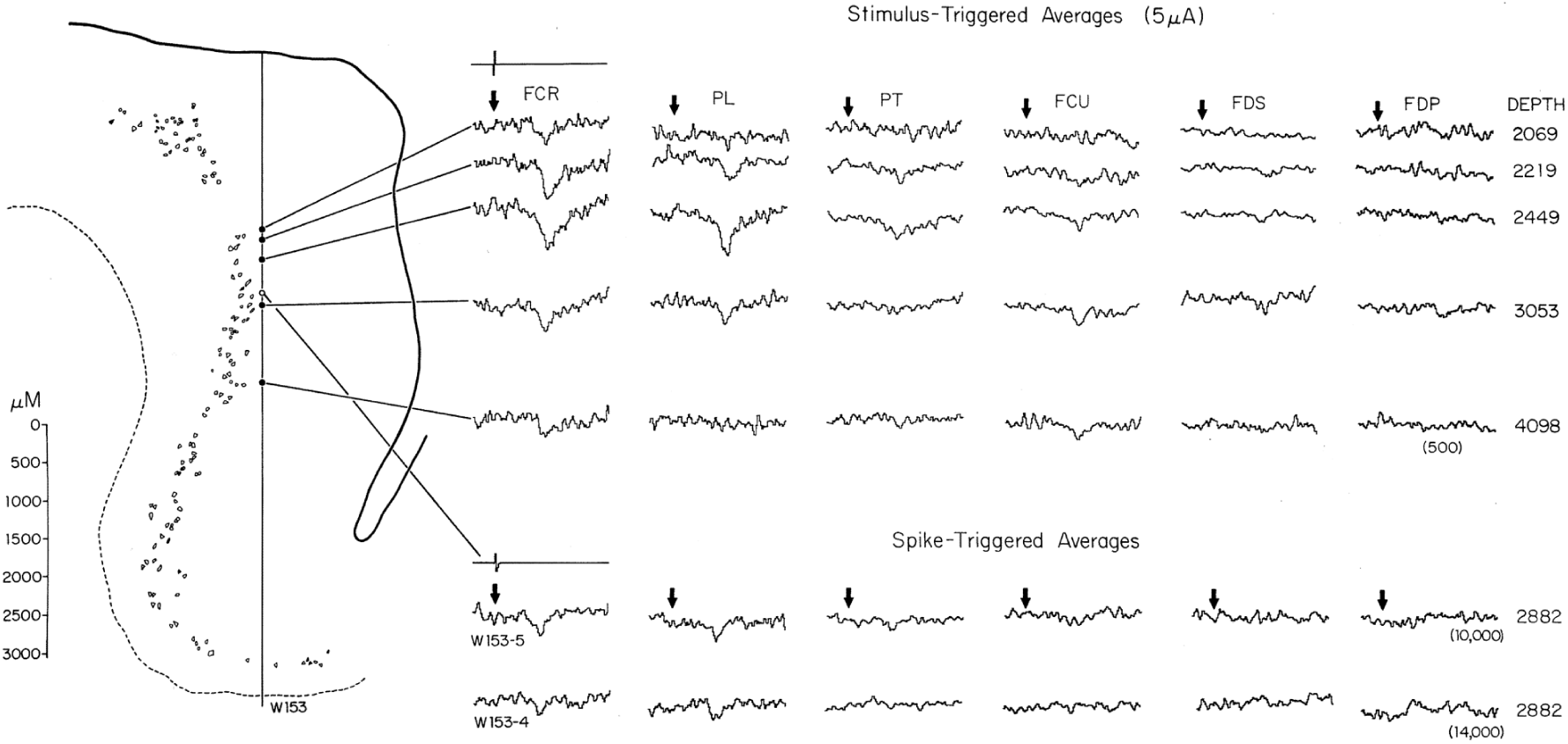
Poststimulus suppression in track W153. Suppression in stimulus-TAs of flexor muscles obtained with 5 μA applied at sequential sites along track W153 matched that seen in spike-TAs triggered from cells W153-4 and W153-5 recorded at depth 2882 (open circle). Layer V cells from Nissl- stained sections are schematically outlined.

**FIG. 6.**
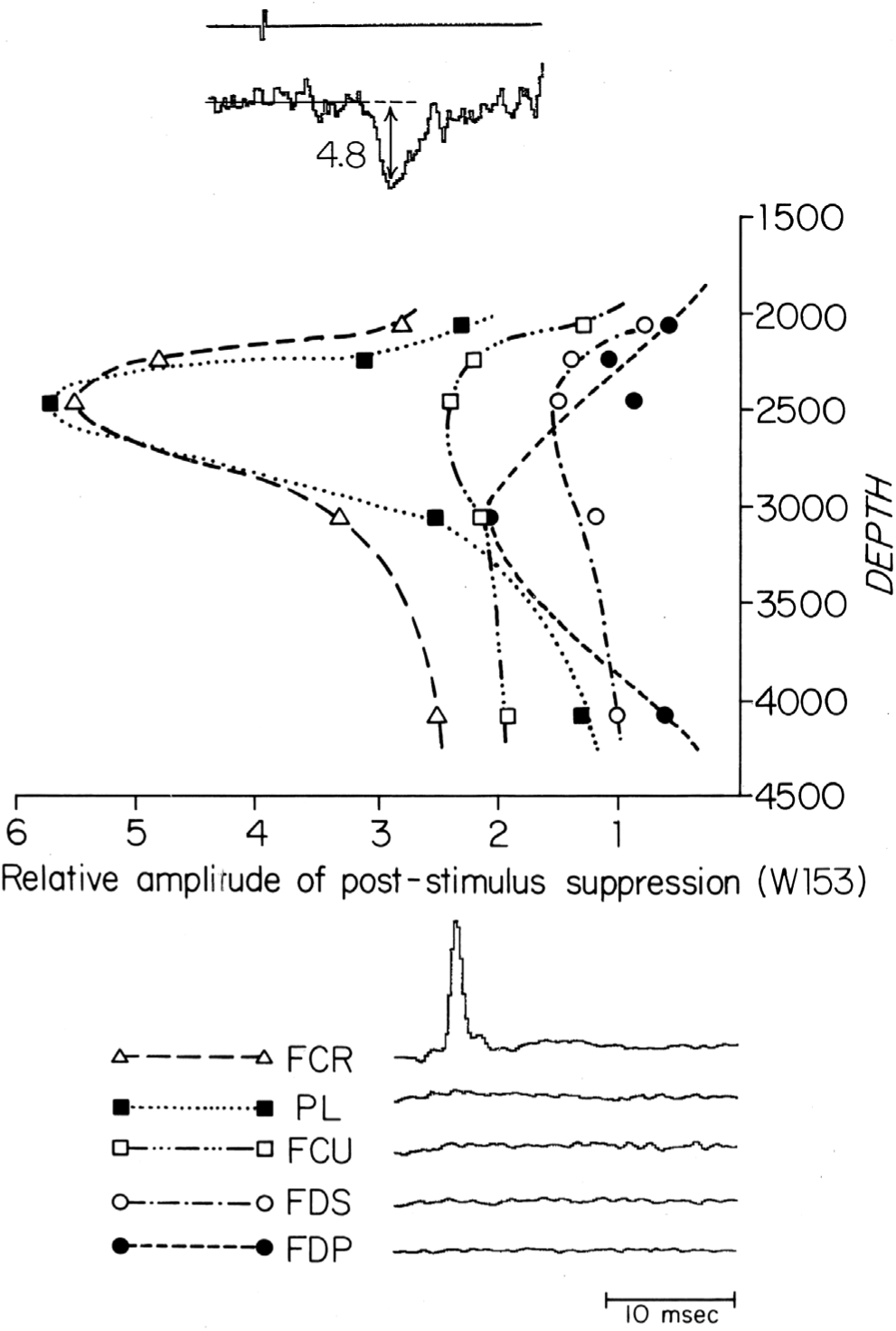
Top: depth-amplitude plot of poststimulus suppression of flexor muscles in track W153 (cf. Fig. 5). Maximal suppression was evoked at depth 2449. Strongest suppression occurred in FCR and PL; lesser effects, with the same distribution over depth, occurred in FCU and FDS. Bottom: EMG-TAs of flexors triggered from FCR motor units showed negligible cross-talk between FCR and the other flexors that were suppressed from microstimuli applied in track W153.

**FIG. 7.**
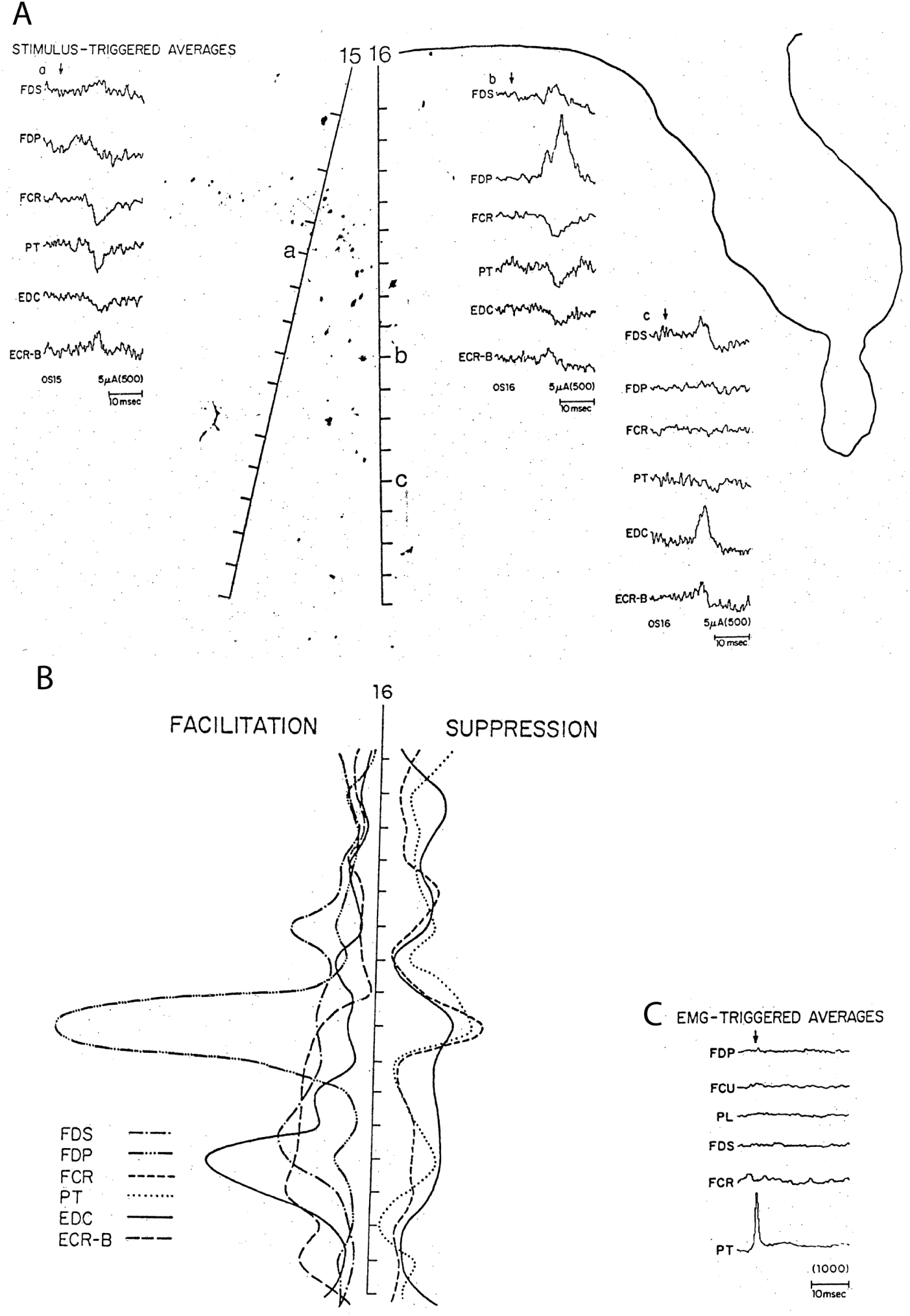
A: stimulus-TAs of muscles evoked from stimulus sites in neighboring tracks OS15 and OS16 with 5-μA stimulation. Stimulation at site 16b produced strong poststimulus facilitation in FDP, but also produced weak poststimulus suppression in FCR and PT equivalent to suppression in FCR and PT evoked from neighboring site 15a (cf. Fig. 11). B: depth-amplitude plot of poststimulus facilitation (to the left) and suppression (to the right) in track OS16. C: EMG-TAs of flexor muscles triggered from motor units of PT confirmed that suppression seen in PT and FCR was not due to crosstalk between these two muscle EMG recordings.

**FIG. 8.**
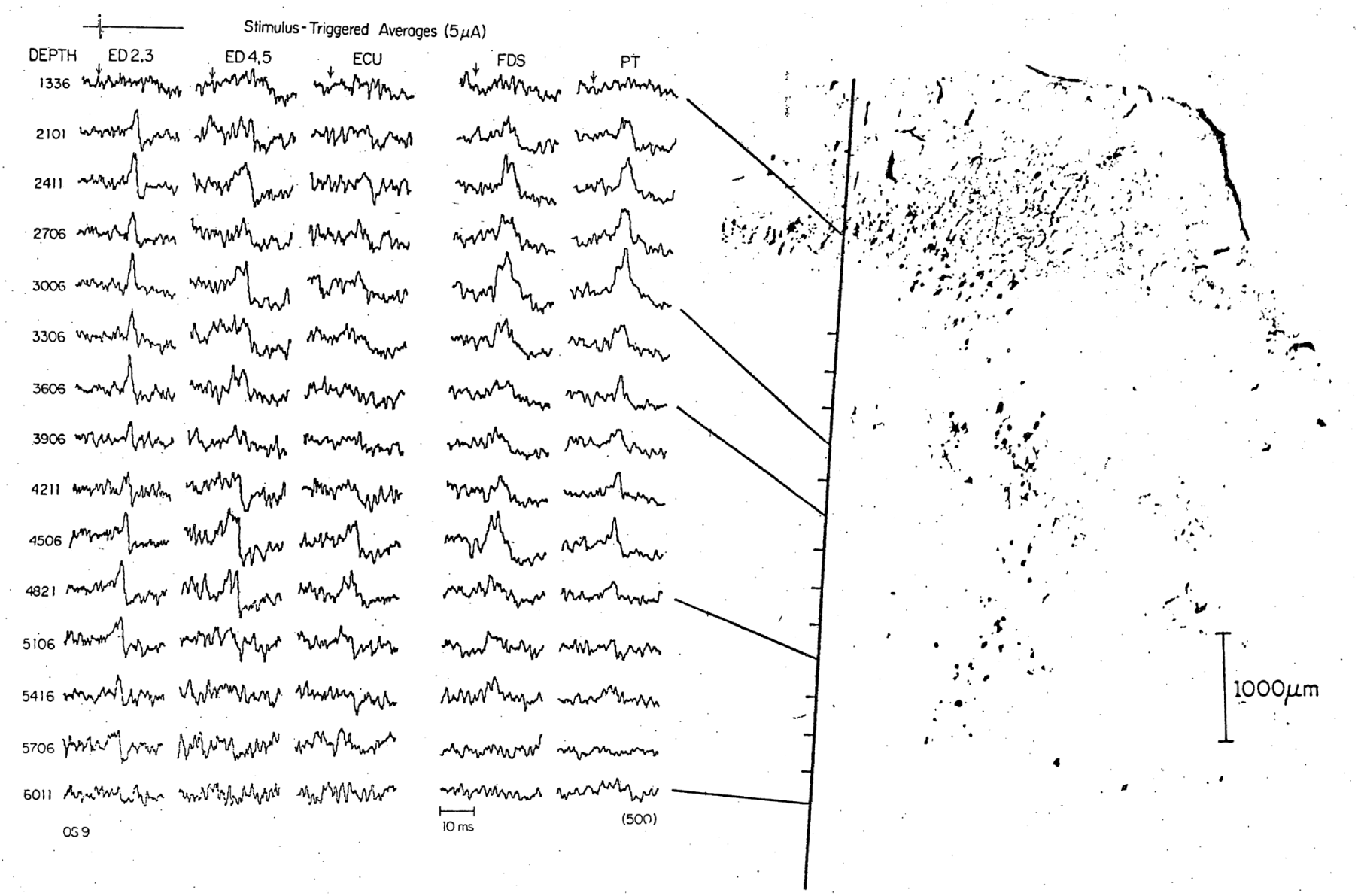
Histologic reconstruction of track OS9 traversing axons in white matter. Track OS9 is superimposed on a parasagittal section (see dashed line in Fig. 9) with HRP-labeled corticospinal cells nearest the track. Corresponding effects in stimulus-TAs of extensor and flexor muscles elicited by 5-uA microstimuli are illustrated. Sites are indicated by a short line to the left of the track and by connecting lines to averages. Biphasic effects were seen in extensors ED2,3, ED4,5, ECU, and ECR-B (not shown). Effects in ECU were complex. Facilitation effects in flexors FDS, PT and FCR (not shown) overlapped with extensor effects, especially at depth 4500 where there was a significant later suppression in extensors.

**FIG. 9.**
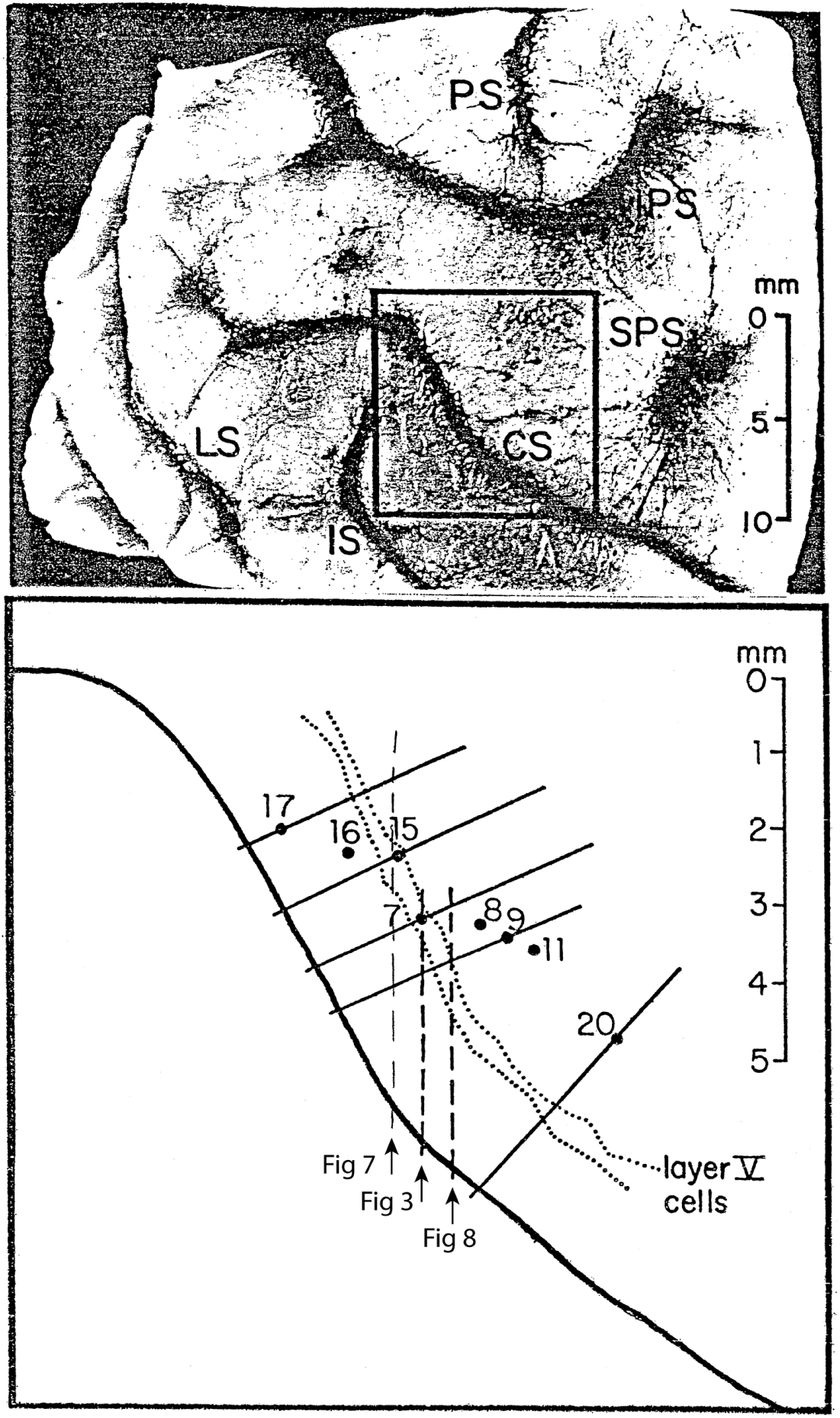
Top: Cortical surface photo of left hemisphere of monkey OS. Bottom: drawing of square area showing location of electrode penetrations (numbered) and layer V cells along the precentral bank (dotted lines). Heavy dashed lines indicate the parasagittal sections shown in Fig. 3 (track OS7) and Fig. 8 (with projected track OS9). The five solid lines indicate the planes of the reconstructions illustrated in Figs. 10 and 11; these reconstructions were compiled from parasagittal sections containing (a) the electrode tracks, (b) the layer V cells closest to the tracks, and (c) the central sulcus contour in the plane of reconstruction. (Track OS16 was superimposed on a reconstruction of track OS15. Similarly, tracks OS8 and OS11 were superimposed on reconstructions of tracks OS7 and OS9, respectively.) CS, central sulcus; IPS, inferior precentral sulcus; IS, intraparietal sulcus; LS, lateral sulcus; PS, principal sulcus; SPS, superior precentral sulcus.

**FIG. 10.**
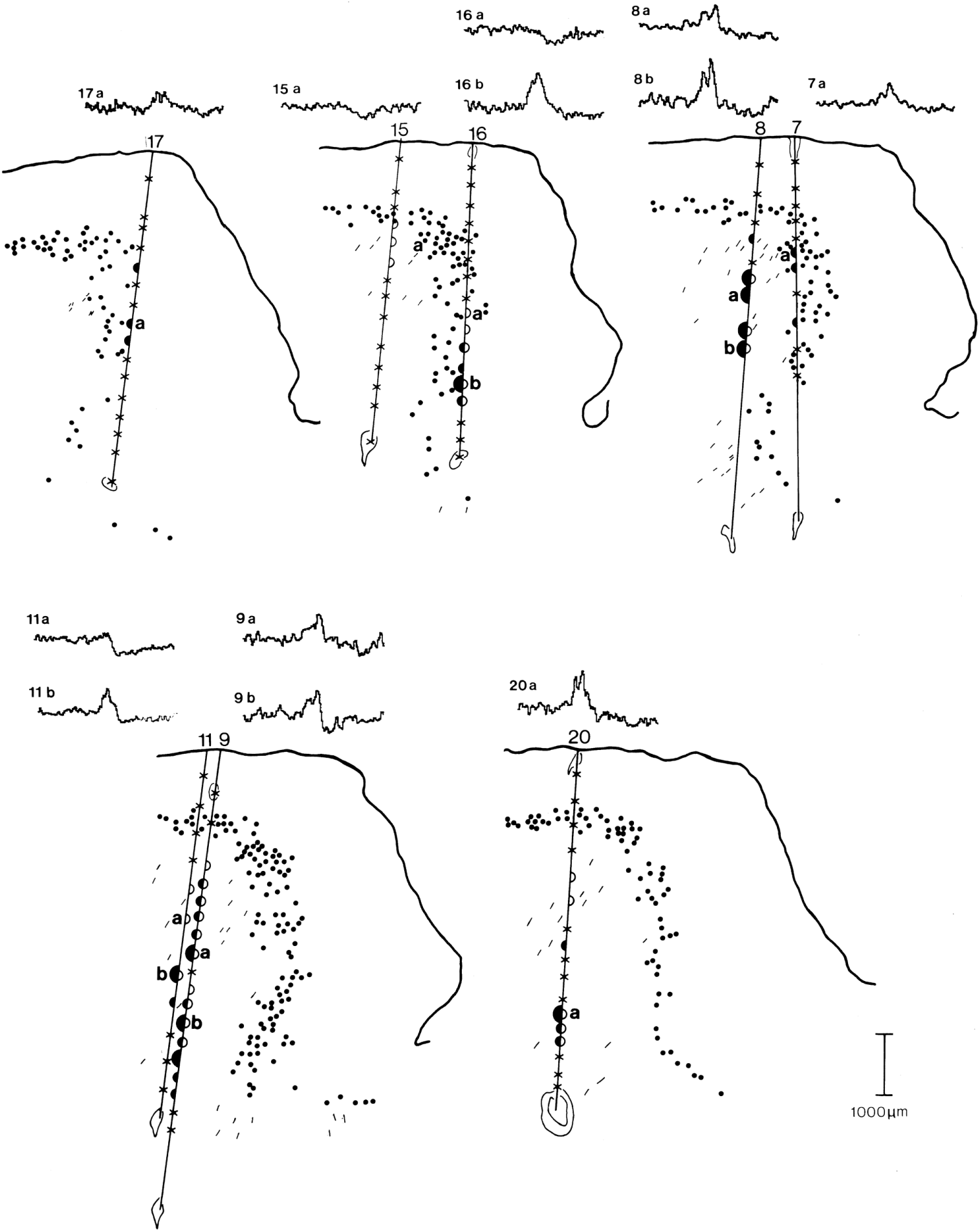
Poststimulus effects evoked in EDC by 5-μA S-ICMS at different cortical sites. Reconstructions (solid lines in Fig. 9) of electrode tracks, layer V cells extending down the precentral bank, and central sulcus contour obtained from parasagittal sections. Large and small half-circles indicate strong and moderate effects, respectively, in EDC evoked from these eight tracks. Solid half-circles to the left indicate facilitation; open half-circles to the right indicate suppression. Xs indicate sites stimulated with no effect in EDC. Lesions and entry points are outlined lightly. Short line segments in white matter indicate HRP-labeled axons. (Note: Track OS9 approximately followed the angle of section. HRP-labeled axons were calculated by trigonometry to project at a 34° angle to the section, and their orientation was at a 20° angle to track OS9.)

**FIG. 11.**
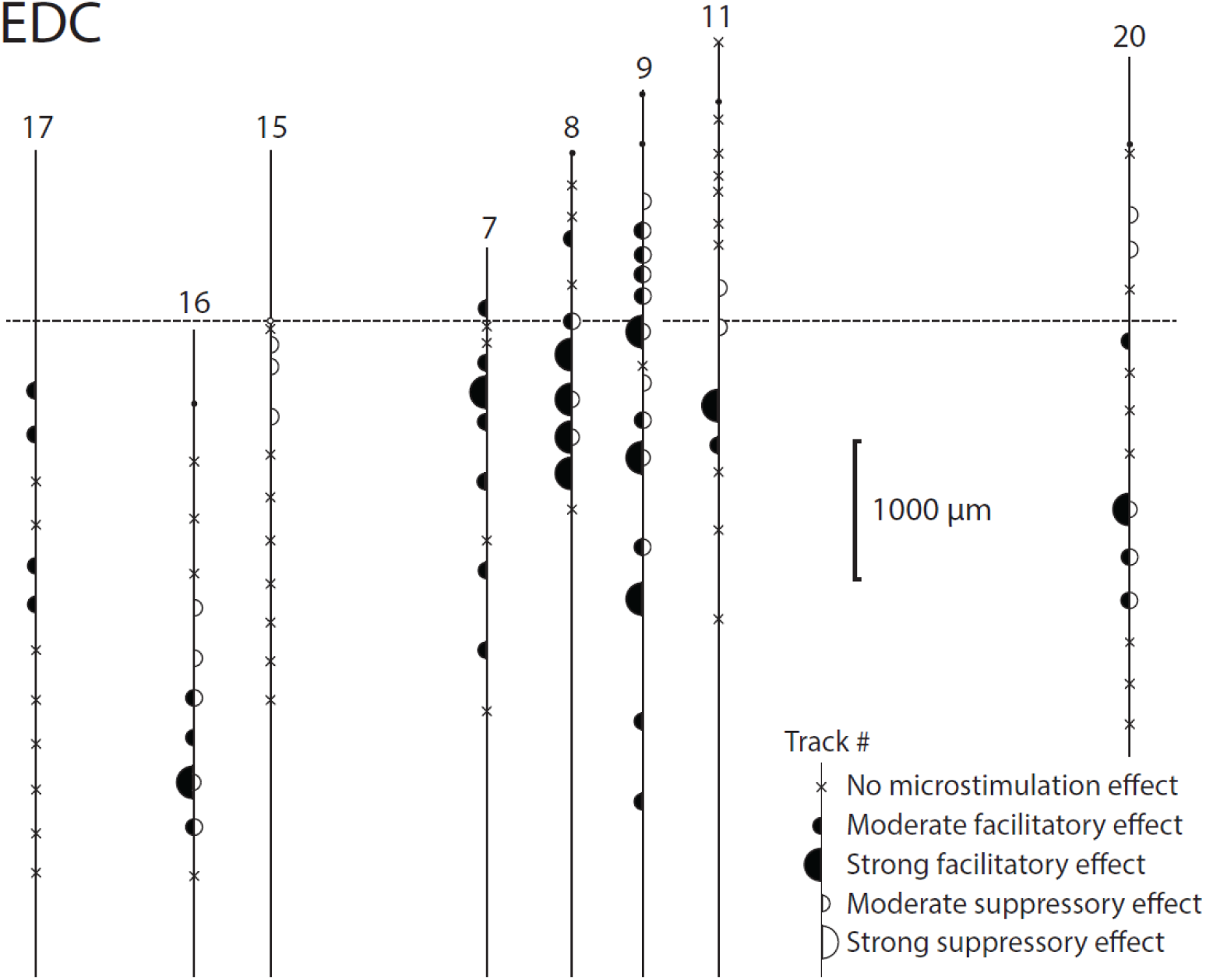
Flattened map of stimulus sites projected to layer V, showing post-stimulus effects evoked in EDC. Dashed line represents location of cortical curvature.

**FIG. 12.**
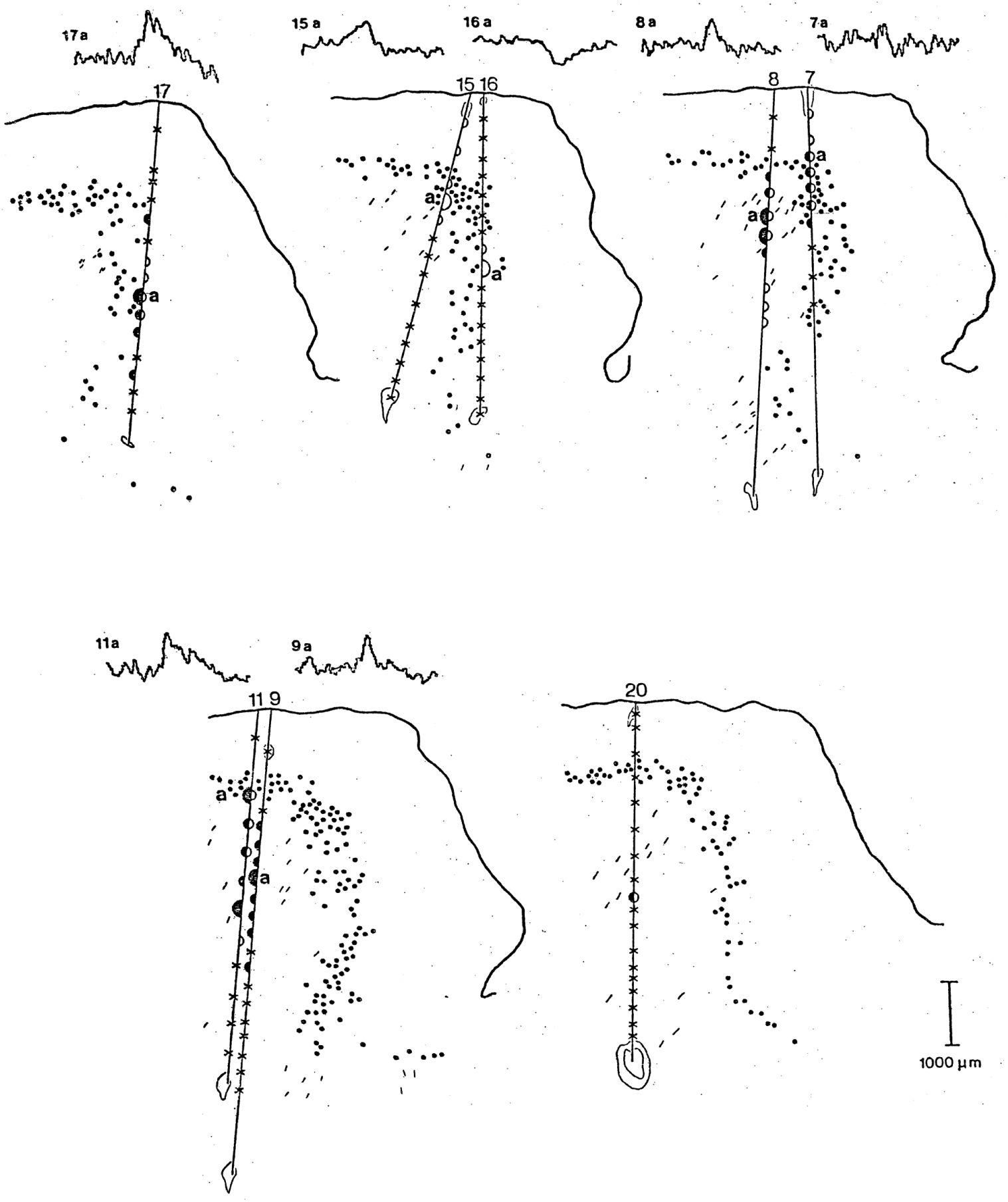
Poststimulus effects in FCR obtained from eight tracks in monkey OS. The tracks and stimulus sites are shown in the reconstructed planes shown by solid lines in Fig. 9. Symbol convention is same as in Fig. 10.

**FIG. 13.**
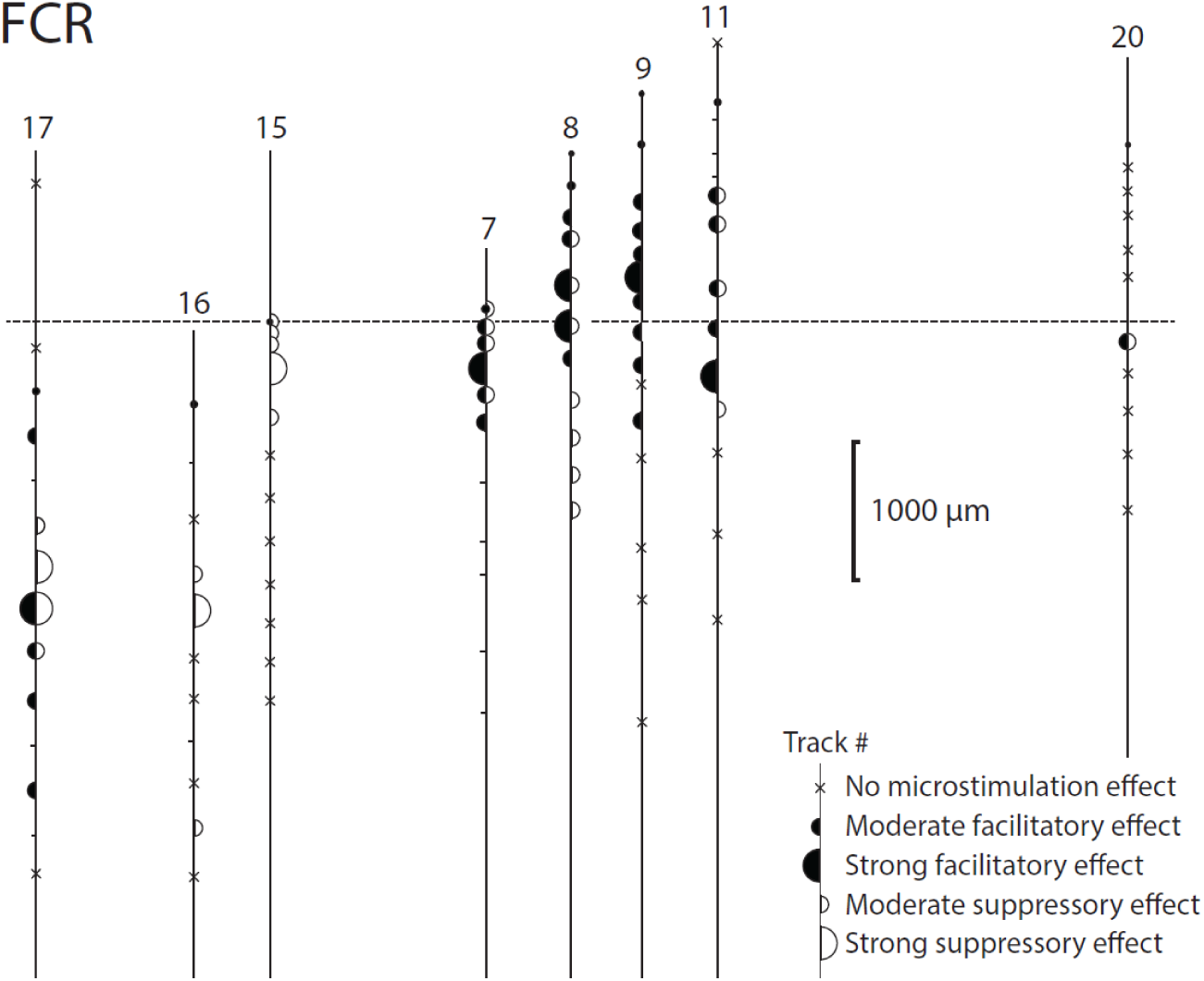
Flattened map for tracks and stimulus sites projected to layer V showing sites with post-stimulus effects in FCR.

## RESULTS

### Extent of output zones producing poststimulus facilitation

Figure 1 illustrates a track in the precentral gyrus which ran tangential to layer V cells, identified in subsequent Nissl histology. This track contained a CM cell (located at symbol o) that produced postspike facilitation in three extensor muscles (ED4,5, EDC and ECR-L). Microstimuli applied at multiple neighboring sites along the precentral bank evoked a similar amplitude profile of poststimulus facilitation in these muscles. Plotting the relative amplitude of poststimulus facilitation versus depth (Fig. 2) revealed peaks in the depth-amplitude plots for the three facilitated muscles around the CM cell; the mean half-width of the peaks was 383 μm. A second, more superficial peak appeared for ECU and ECR-L. EMG-TAs confirmed that facilitation in multiple muscles was not just the result of common recording of the same motor units with different intramuscular electrode pairs; for example, Fig. 2 (bottom) illustrates the absence of cross-talk from ECR-L to the other three muscles.

In some tracks microstimuli facilitated both flexor and extensor muscles, although the maximal effects in each group were usually evoked at separate sites. In Fig. 3, poststimulus facilitation in FDS was maximal for microstimuli applied at depth 1600, while EDC was activated maximally at depth 2219, and again at depth 3514. The depth-amplitude plots of poststimulus facilitation in FDS, PL, EDC and ECR-L are drawn in Fig. 4A; the dashed lines indicate the distances over which effects were highly significant (P < 0.01). There was no overlap of three statistically defined output zones -- a superficial zone facilitating FDS and PL, an intermediate zone facilitating EDC and ECR-L, and a deeper zone facilitating just EDC. Histological reconstruction of this track (Fig. 3) indicates three clusters of HRP-labeled corticospinal cells corresponding to these zones.

The distances over which poststimulus effects were statistically significant (P < 0.01) and the half-widths of output zones were tabulated for those electrode tracks that passed tangentially near layer V cells extending down the bank of the precentral gyrus. The mean half-width of six facilitation output zones was 750 + 50 μm (SEM), for S-ICMS of 5-μA (n = 11 muscles). For 10-μA S-ICMS the mean half-width of two other zones was larger: 1100 + 100 μm (n = 8). The mean distances over which S-ICMS evoked statistically significant poststimulus effects (P < 0.01) were 950 + 50 μm at 5 μA, and 1400 + 100 μm at 10 μA. The half-width measurement was less than the P < 0.01 level and may have underestimated the size of output zones.

As described previously (8), microstimuli applied at some precentral sites affected both flexor and extensor muscles. Sites which activated one set of muscles (called agonists) evoked suppression, or facilitation, or no effect on the antagonist muscles, with approximately equal probability. Similar combinations of effects were observed for the output zones in this study. Microstimulation in track W125 which facilitated extensor muscles (Fig. 1), also suppressed FCU and PL at the deeper zone. The track represented in Figure 4 contained sites which facilitated muscles without suppressing their antagonists. For example, at depth 1600, stimuli facilitated flexors but produced no significant effect in the antagonist extensor muscles.

### Extent of output zones producing poststimulus suppression

S-ICMS sometimes elicited pure suppression without any effect in the antagonists of the inhibited muscles. Fig. 5 illustrates such a track, which also contained two cells that produced a postspike suppression of the same muscles (6). Cells W153-4 and W153-5 were active during flexion but not during extension, and, somewhat counter-intuitively, spike-TAs revealed postspike suppression of two muscles active during flexion: FCR and PL. S-ICMS with 5-uA current pulses applied in the vicinity of these cells evoked poststimulus suppression in FCR and PL. Applied during extension S-ICMS produced no effects in the recorded extensor muscles. Depth-amplitude plots (Fig. 6 Top) of the poststimulus suppression indicated that FCR and PL were significantly suppressed over distances of about 1000 μm. The mean tangential length of output zones along the precentral bank based on statistically highly significant (P < 0.01) poststimulus suppression was 950 + 100 μm, n = 6 (Table 2). For six cases of poststimulus suppression evoked by 5-uA S-ICMS, the mean half-width of the suppression zones on the precentral bank was 850 + 100 μm (two tracks in two monkeys, Table 2).

### Mixed effects in agonists from microstimulation at single cortical sites

At some cortical sites S-ICMS elicited post-stimulus facilitation mixed with suppression in synergistic muscles. Figure 7 illustrates such mixed post-stimulus effects evoked in six muscles from representative sites. Microstimuli applied at site 16b of track OS16 strongly facilitated FDP, while producing clear poststimulus suppression of FCR, PT and EDC. (The orientation of this track and the relative location of HRP-labeled corticospinal cells on the precentral bank are shown in figures 10 and 11.) The depth profile of the post-spike effects in these muscles reveals two distinct zones. Mixed effects evoked in synergists from a single site could be due to activation of cells belonging to different output zones. No CM cells have yet been found to produce such mixed post-spike effects in synergist muscles.

### Mapping sites for evoking facilitation or suppression of the same muscles

To map the output sites more systematically, a series of tracks were made parallel to the precentral bank in monkey “OS” whose corticospinal cells were subsequently labelled with HRP. A surface view of the cortex of this monkey (Fig. 9B) shows the entry points of the electrode tracks (numbered), parasagittal histological sections shown in Figs. 7, 3, 8 (dashed lines), five reconstructed sections (solid lines) shown in Figs. 4, 10, 11 and the projected locations of layer V cells lying along the precentral bank (dotted lines). The solid lines represent the reconstruction planes which contain specific electrode tracks and are oriented perpendicular to the plane of the layer V cells (and therefore contain the corticospinal cells closest to the tracks). The stimulation sites and the layer V cells in these planes were reconstructed from the closest parasagittal histologic sections (oriented parallel to the dashed lines).

To document the sites affecting a single muscle Fig. 10 illustrates the location of output sites affecting the EDC muscle, and the nearest HRP-labeled corticospinal cells in these planes. Stimuli at effective sites evoked poststimulus facilitation (filled semicircles) or suppression (open semicircles) or both. Representative averages are also shown, to illustrate these effects. EDC was facilitated from multiple sites along the precentral bank (from tracks OS17 to OS20). The dorsoventral dimension of the region containing sites which facilitated EDC was 4.5 mm. Suppression of EDC was obtained from a region 4.5 x 3 mm (mediolateral x dorsoventral, from tracks OS16 to OS20).

The tracks illustrated in Fig. 10 contain many stimulus sites in white matter, which contained axons of cells whose soma resided in layer 5. To represent the two-dimensional spatial distribution of the output zones in layer V more clearly, the stimulus sites in white matter were projected to corresponding layer V cells by following the axonal trajectories defined by HRP-labelled fiber segments. The surface defined by layer V was then unfolded onto a plane. The resulting map for EDC is shown in Fig. 11.

The projected map of stimulus sites for FCR is shown in Fig. 12 and the flattened map for FCR is shown in Fig. 13, indicating the relative location of the layer V cells presumed to have mediated the output effects on FCR. The resulting grid of stimulus sites is not dense enough to fully define the contours of the output zones, but the sampled sites indicate that zones for individual muscles have dimensions of several hundred microns.

Furthermore, there is some indication that excitatory bands are interspersed with inhibitory regions. For FCR this alternating pattern is shown by the inhibition in tracks 16 and 17, interspersed between neighboring excitatory tracks. The flattened map for ECR-B is shown in Fig. 14. Pure inhibition was evoked from many sites in track 9.

**FIG. 14.**
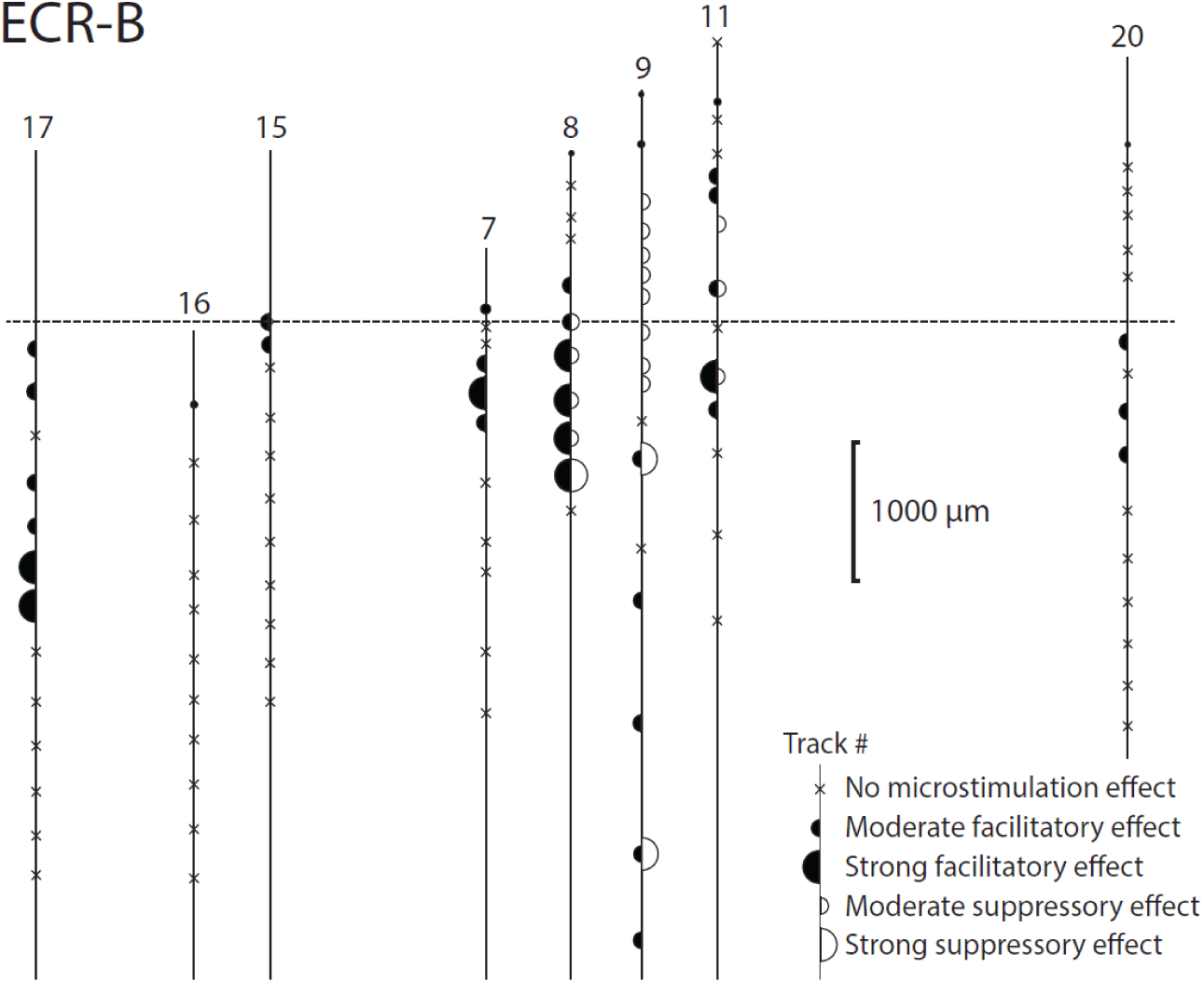
Flattened map for muscle ECR-B.

These purely inhibitory effects were in contrast to suppression following excitation, which could also be due to the post-firing refractory period of motor units. Comparing Figs 13 and 14 shows that the two antagonist muscles, FCR and ECR-B, were reciprocally affected from some sites (e.g., tracks 9 and 15), but they were also coactivated from other sites (e.g., in tracks 7 and 17).

The illustrated results are representative of the distribution of output sites for other muscles. Flattened maps for two additional muscles, PL and FDP are shown in supplemental figures S1 and S2. Comparing these maps reveals that sites that facilitated flexors were often intermingled with those that facilitated extensors.

### Cortical distribution of single CM cells facilitating forelimb muscles

To characterize the spatial distribution of single CM cells facilitating individual muscles, as seen in the dorsal view of the 2D plane of the recording chamber, we mapped the locations of CM cells producing clear postspike facilitation in each muscle. The CM cell distributions for seven forearm muscles in monkey W are illustrated in Fig. 15. The regions containing CM cells which facilitated a particular muscle typically overlapped the loci of CM cells affecting other muscles. The mediolateral and dorsoventral dimensions of regions containing CM cells affecting a given muscle ranged from 3 to 7 mm. For example, ED2,3 showed facilitation from cells in different tracks extending from sites at depths 753 to 4732, a dorsoventral dimension of about 4 mm.

**FIG. 15.**
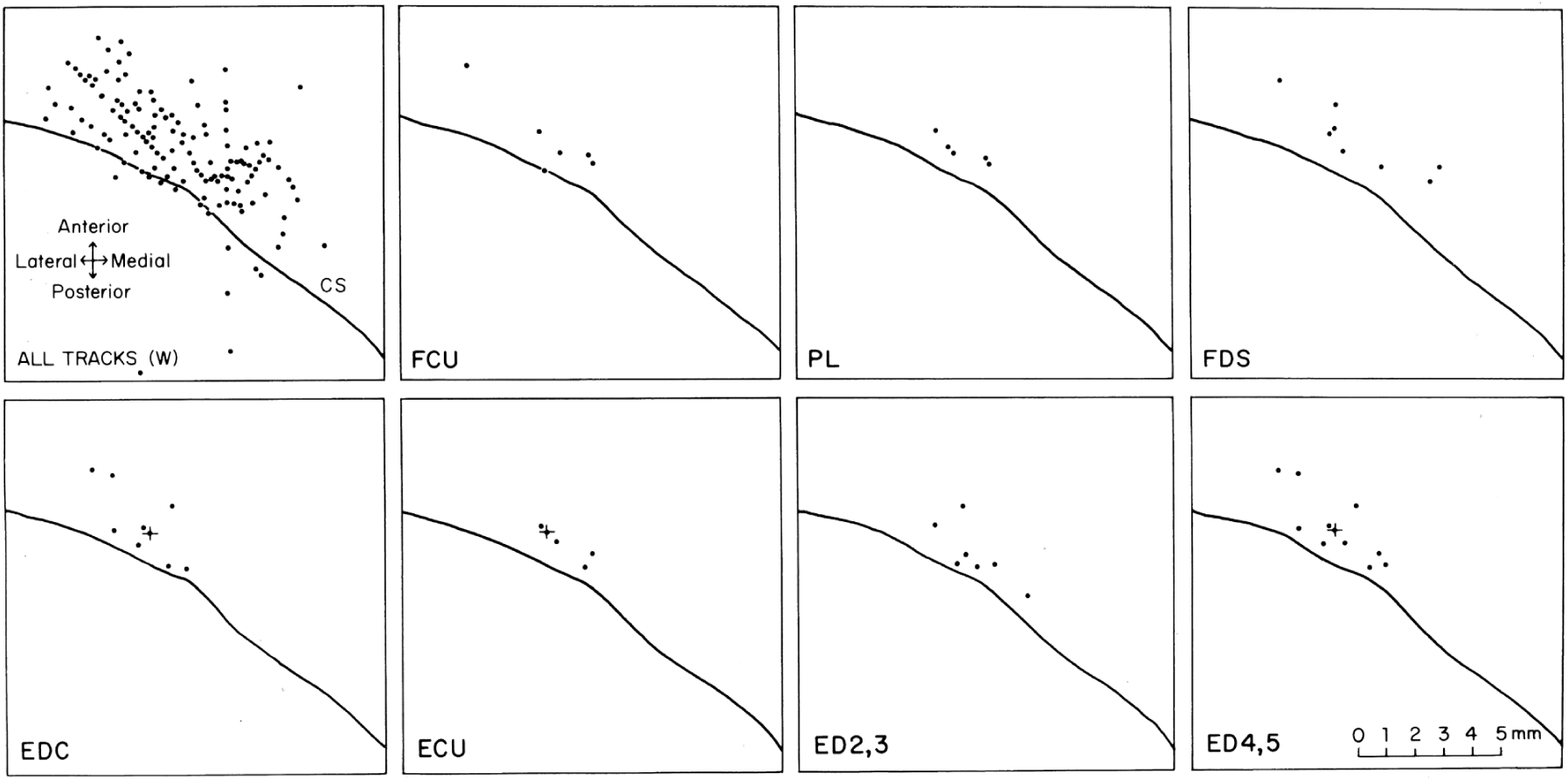
Spatial distribution of CM cells with moderate or strong postspike facilitation in particular muscles. Dots represent dorsal view of entry points of 132 tracks made in this monkey (W). Position of central sulcus was determined from histology. Many CM cells had muscle fields including several agonists. + shows a CM cell with postspike facilitation in EDC, ECU, ED4,5, ECR-B and ECR-L. There are 26 different CM “muscle fields” represented here. Note the overlap of regions with CM cells affecting different muscles.

**FIG. 16.**
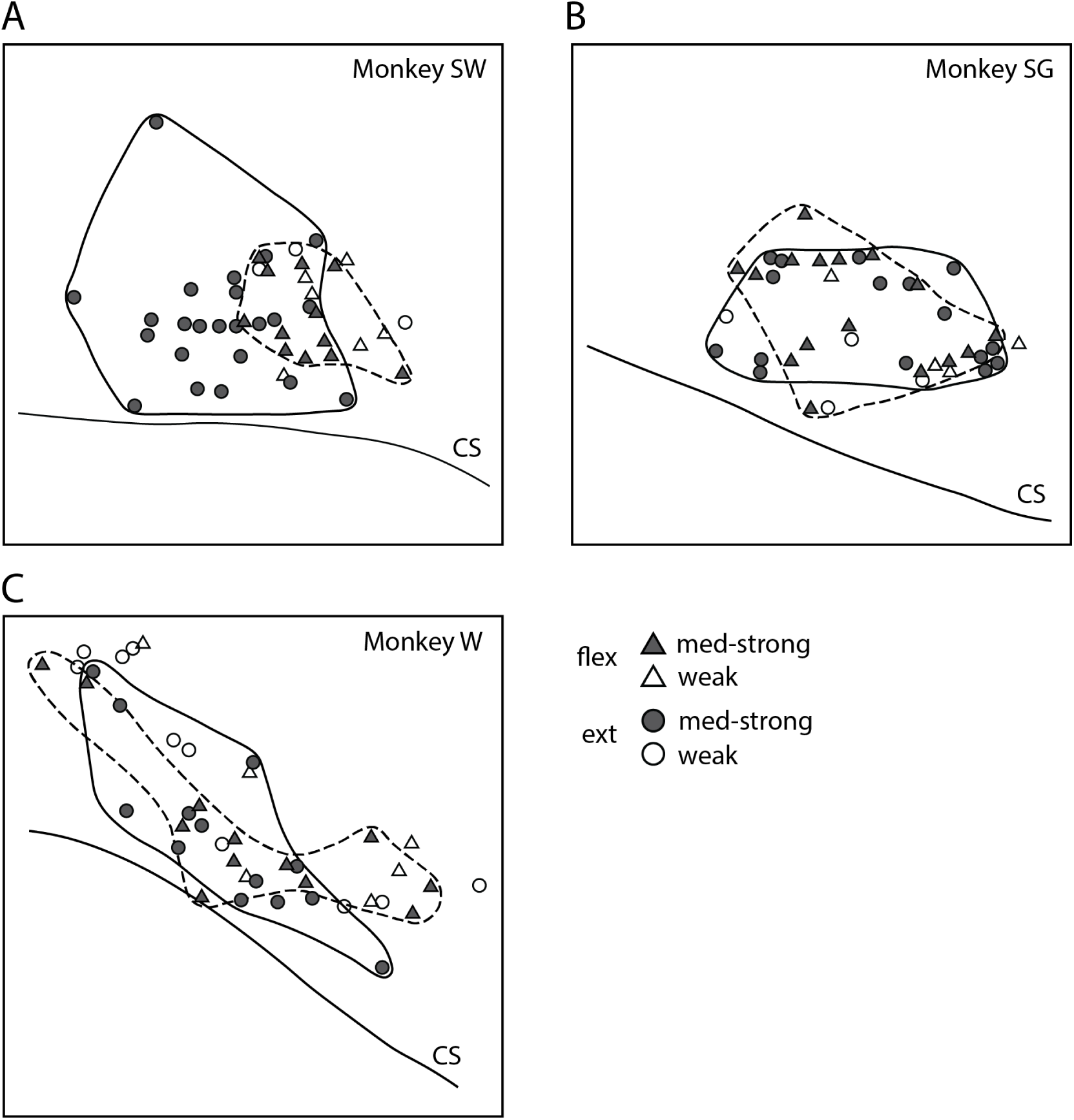
Regions containing CM cells that facilitated extensor and flexor wrist and digit muscles for three different monkeys. Triangles and circles represent tracks with CM cells facilitating flexor and extensor muscles, respectively. Filled symbols represent moderate to strong facilitation effects. Open symbols represent weak effects. Regions with CM cells having moderate to strong effects are circled with a dashed line (flexion CM cells) or a solid line (extension CM cells). Monkeys W and SG had overlapping flexion and extension regions; monkey SW had many CM cells that facilitated extensor muscles in a region without any CM cells that facilitated flexor muscles.

Since many CM cells produced postspike facilitation in more than one target muscle, it was of interest to examine the cortical distribution of muscle fields. In this monkey the CM cells exhibited 26 different muscle fields; 21 of these were represented in 21 different electrode penetrations. Only 5 of the 26 muscle fields were replicated in more than one electrode track, i.e., appeared in different output zones.

The region on the precentral bank that contains CM cells facilitating flexor muscles of the wrist and digit and the region that contains extensor CM cells were determined in three monkeys. In two monkeys these two regions overlapped, while in the third they were largely separate (Fig. 15). The mediolateral dimension of the regions that affected all agonists was similar to that of regions that affected only one muscle; this distance ranged from 2 to 7 mm in the three monkeys.

## DISCUSSION

### Dimensions of cortical output zones along the bank of the precentral gyrus

This study presents new data on the extent of motor cortical output zones affecting multiple wrist and digit muscles in primates. This study combines the finest level of controlled activation (using subthreshold single-pulse ICMS, usually 5 μa) with detection of subthreshold excitatory and inhibitory effects on muscles (using stimulus-triggered averages). The locations of cortical stimulus sites were also defined in relation to HRP labelled corticospinal neurons and the locations of single CM cells (which had postspike effects on the same muscles). Our results support the view that precentral cortex contains clusters of cells or zones in which each cell of the cluster or zone produces a similar profile of output effects (muscle fields or muscle synergies). These regions often contain CM cells that facilitate the same target muscles, but they clearly extend beyond the CM cells.

Similar muscle synergies were seen in stimulus-TAs over distances of 500-1250 μm when stimuli were delivered along electrode penetrations aligned tangential to layer V cells in the precentral bank. Depth - amplitude plots showed that the amplitude of poststimulus facilitation or suppression in a set of muscles typically increased and decreased together.

The extent of these output zones could not be attributed entirely to current spread. Studies of current spread from S-ICMS in cat motor cortex indicated that 5 μA activated cells directly within a radius of 65 μm, while 10 μA spread 90 μm (32). Larger estimates of physical current spread have been obtained for spinal interneurons, namely, 80-100 μm for a 5-uA stimulus (16). Indirect, transsynaptic activation of cells is effective at distances of at least 125 μm from the site of repetitive stimulation at 5 μA (16, 32); transsynaptic excitation may also have contributed to spread with our single pulse ICMS. In comparison, the dimensions of specific poststimulus effects we observed were 500-1250 μm for 5-μA current and 800-1850 μm for 10 μm current (P < 0.01).

To determine the minimal dorsoventral dimension of our cortical output zones, we subtracted 150 μm from the measured extent to account for physical spread of a 5-uA stimulus. The resulting average dorsoventral dimension of cortical output zones of both poststimulus facilitation and suppression (P < 0.01) on the precentral bank was then 800 μm.

These dimensions are comparable to those obtained by repetitive ICMS; for example, Asanuma and Rosen (3) found radially oriented output “columns” that had diameters of about 1 mm. They are also consistent with reports of Kwan et al (21) that discrete movements (e.g., extension of digits 2-5) could be evoked from dorsoventral distances of 0.5-1.0 mm along the bank of the precentral gyrus. These dimensions are also comparable to those in 2D maps of 24 individual arm muscles (including shoulder, elbow, wrist and digit muscles) obtained from systematic mapping of the precentral cortex with 15 μA stimulus triggered averaging during reaching movements (16, 28).

Our results also agree with the anatomical findings of Jones and Wise (19), that HRP-labeled corticospinal “clusters” had widths of 0.5-1.0 mm in parasagittal sections. The HRP-labeled cells in our sections also seemed to be grouped in clusters of about 1000 μm in diameter (Figs. 3 and 8). In contrast, Murray and Coulter (26) found that HRP-labeled corticospinal cell clusters were much less distinguishable in cortical areas with high cell density such as the precentral forelimb region than in less dense areas.

### Extent of cortical output sites for particular muscles

The mediolateral dimensions of regions containing all the output zones affecting a particular muscle ranged from 4.5 to 5 mm (e.g., Fig. 10 for EDC). The dorsoventral dimensions of these regions ranged from 3.5 to 4.5 mm (Fig. 10). The mediolateral dimension is comparable to data on regions pooled from several monkeys containing discontinuous “efferent zones” having a specific effect (e.g., FDS, 4 mm mediolateral; digit 2,3 extension, 8 mm mediolateral) (2). Like Asanuma and Rosen (3), Kwan et al. (21) and Hudson et al. (16) we found substantial overlap between regions affecting wrist and digit muscles and antagonist muscles; the output zones to a specific muscle overlapped with regions affecting synergists and antagonists. Andersen and coworkers (1) found that cortical areas where repetitive ICMS elicited effects in a single muscle were as large as 6 x 5.5 mm. This difference with our results could be attributed to spread of effects via temporal summation of stimulus trains (17). They also found overlap between areas affecting EDC, thenar muscle, and first dorsal interosseus.

Our results agree with those of Kwan et al. (21), who showed that with repetitive ICMS (300 Hz) the area of overlapping wrist and digit effects was predominantly on the precentral bank and had mediolateral dimensions of 7 and 5 mm, respectively. In one monkey we found that trains of repetitive ICMS applied within the wrist and digit area evoked movements of these joints only. Trains applied just outside this region evoked biceps contraction on the medial side and pectoralis contraction on the lateral side, consistent with the view that surrounding areas affect elbow and shoulder muscles (21). Usually, the cortical regions that contained output zones affecting both agonist and antagonist muscles of the wrist and digits overlapped.

### Cortically evoked inhibition

As in previous studies with spike- and stimulus-TAs of EMG activity (7, 8, 16, 20, 28), we found many sites which evoked suppression of muscle activity, an effect missed by studies documenting only evoked activation. Stimulation at some sites evoked “pure” postspike suppression without any mixture of facilitation; other sites evoked facilitation followed by suppression in average muscle activity. Interpretation of the latter cases remains ambiguous. Some of the post-peak suppression may be due to the refractory period of the motor units which fired in the peak. If the peak is produced by advancing motoneuron spikes that would have occurred later, the area of the peak should equal the area of a subsequent trough. This has been seen in cases of poststimulus and postspike motor unit histograms (27). In some cases, the trough may be filled in by later distributed excitation. As argued by Lemon, et al (23), the post-peak trough may also reflect genuine post-synaptic inhibition, particularly when a stimulus evokes a potentially mixed volley. Indeed, the corticospinal EPSPs evoked by S-ICMS are often followed by IPSPs. In light of this ambiguity, we have not interpreted the post-peak suppression as unequivocal evidence for inhibition. Nevertheless, it is clear from the plots that pure inhibitory sites were intermingled with excitatory sites. For many muscles the maps reveal pure inhibitory zones interspersed with excitatory regions (Figs. 10, 11).

The inhibitory regions for some muscles were coextensive with excitatory regions for antagonists. This pattern agrees with the finding that some CM cells produce reciprocal inhibition of antagonists of their target muscles (20).

### Distribution of CM cells

The finest level of spatial resolution of cortical output comes from the postspike effects of single CM cells. CM cells affecting a given forelimb muscle were distributed over regions having mediolateral dimensions of 3 to 7 mm. However, CM cells facilitating a given target muscle usually had different muscle fields. In one monkey a particular muscle field was represented redundantly, only 19% of the time (i.e., 21 of 26 distinct muscle fields were observed only once). It seems likely that sampling additional muscles could have shown the muscle fields of these cells to differ in the representation of other muscles. The CM cells affecting a particular muscle were intermingled with CM cells affecting synergists and antagonists.

### Summary and conclusions

Our experiments used the finest level of controlled activation (subthreshold single-pulse ICMS, usually 5 μa) and detection of statistical effects on muscles (stimulus-triggered averages). The results support our previous study showing that the profile of effects in low intensity stimulus triggered averages generally closely resemble the effects in spike-triggered averages obtained from single CM cells at the same site (7, 8). Our results suggest that the motor output regions in the precentral gyrus examined with these techniques consist of zones that affect specific muscles in different combinations. These zones typically contain CM cells with similar distribution of postspike effects in these muscles. Examining the profile of excitatory and inhibitory effects on groups of muscles reveals complex combinations, with little if any redundancy.

## ACKNOWLEDGMENTS

We thank Mr. Jerrold Maddocks and Mr. Larry Shupe for technical help, and Ms. Kate Schmitt for editorial assistance. Supported by National Institutes of Health grants NS12542, RR00166, NS5082 and US0966.

Present address for P.D. Cheney: Emeritus Professor, Department of Cell Biology and Physiology, University of Kansas Medical Center, Kansas City, KS 66103. Present address for S. Sawyer Vincent (was S. Palmer 1983-1993): retired from Dept. of Biology and Chemistry, Oral Roberts University, Tulsa, OK. Present address for R.F. Martin unknown.

## Supplementary Information

**FIG. S1.**
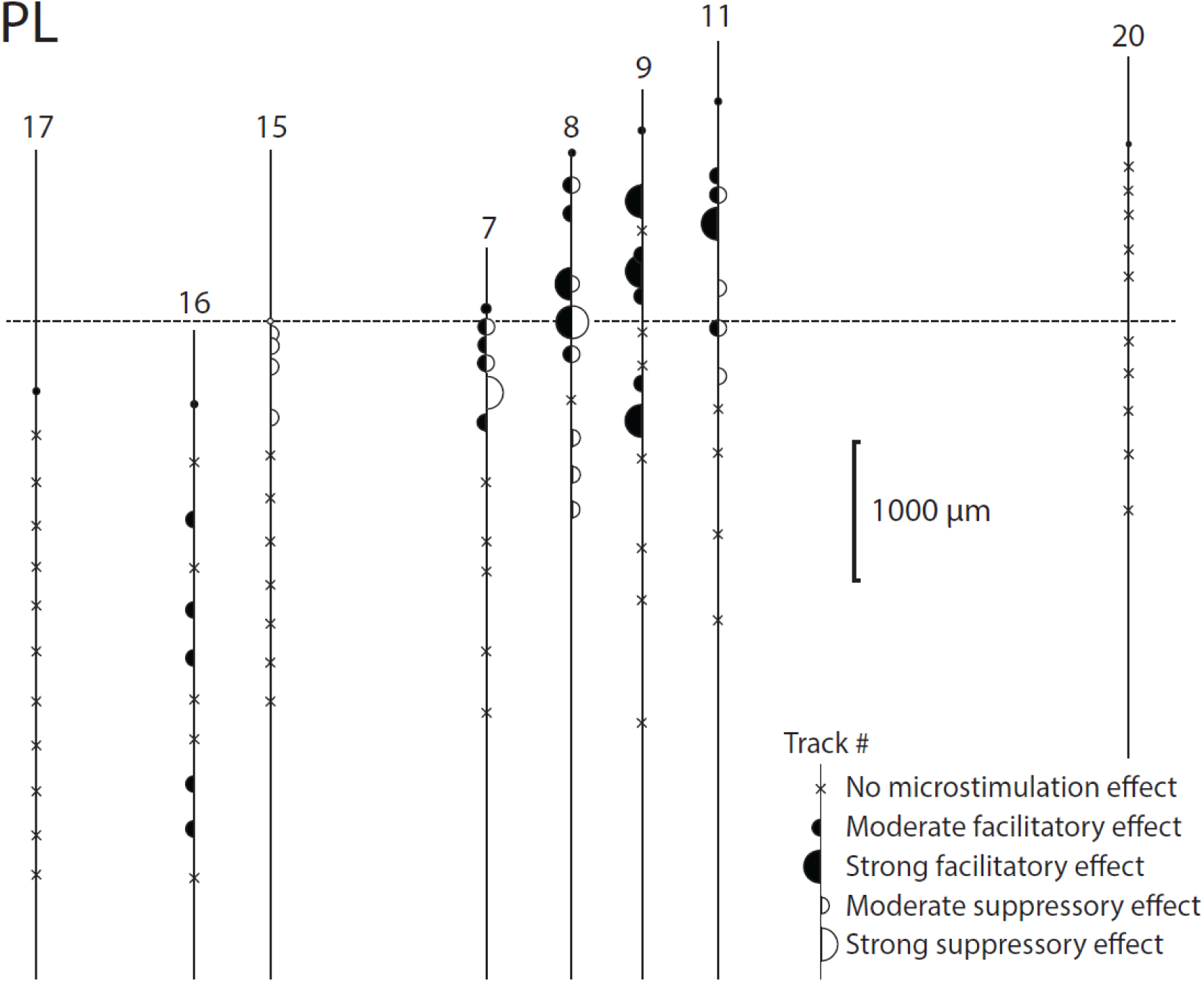
Flattened map of poststimulus effects in muscle PL.

**FIG. S2.**
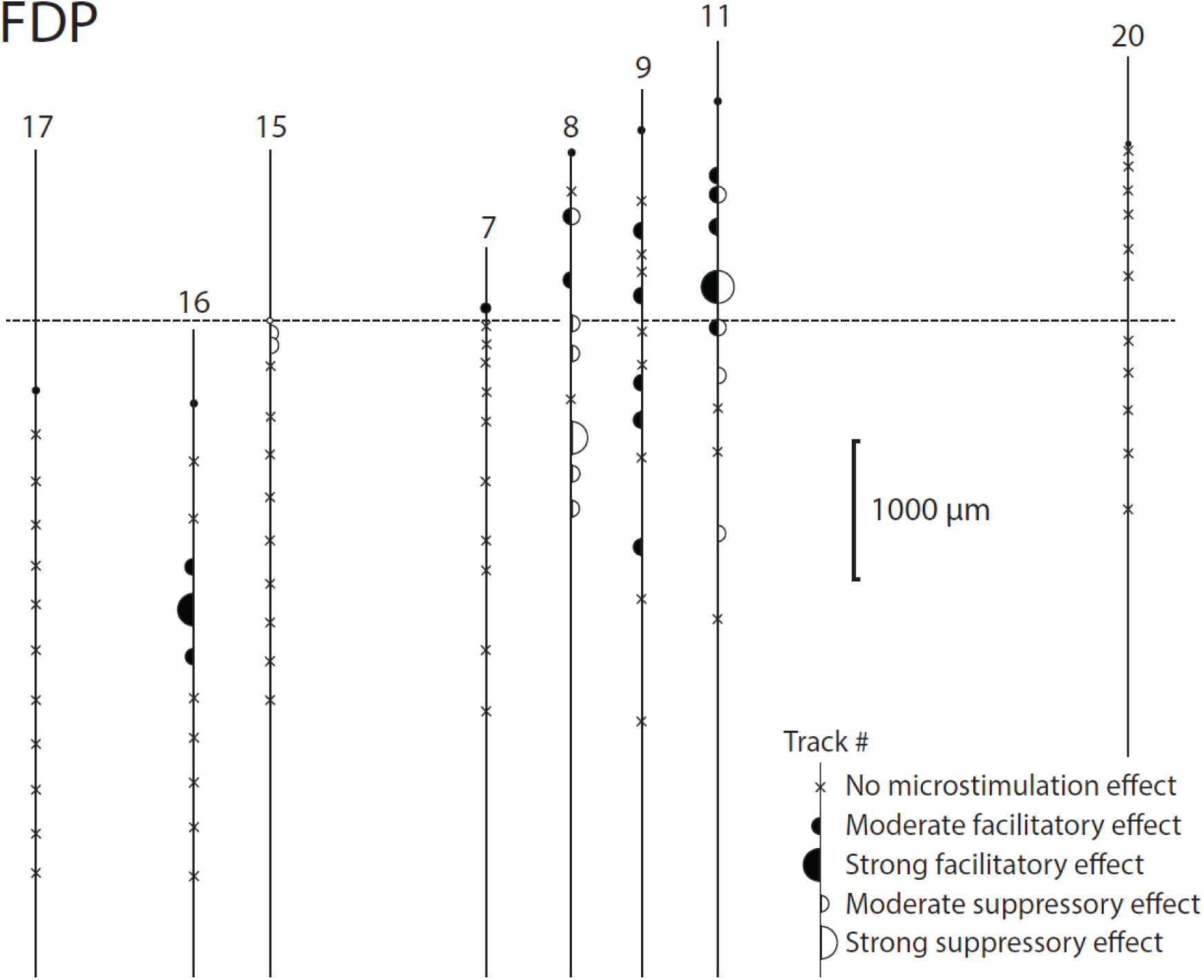
Flattened map of poststimulus effects in muscle FDP.

